# Performance comparison of reverse transcriptases for single-cell studies

**DOI:** 10.1101/629097

**Authors:** Zucha Daniel, Androvic Peter, Kubista Mikael, Valihrach Lukas

## Abstract

**Background:** Recent technical advances allowing quantification of RNA from single cells are revolutionizing biology and medicine. Currently, almost all single-cell transcriptomic protocols rely on conversion of RNA to cDNA by reverse transcription (RT). However, RT is recognized as highly limiting step due to its inherent variability and suboptimal sensitivity, especially at minute amounts of RNA. Primary factor influencing RT outcome is reverse transcriptase (RTase). Recently, several new RTases with potential to decrease the loss of information during RT have been developed, but the thorough assessment of their performance is missing.

**Methods:** We have compared the performance of 11 RTases in RT-qPCR on single-cell and 100-cell bulk templates using two priming strategies: conventional mixture of random hexamers with oligo(dT)s and reduced concentration of oligo(dT)s mimicking common single-cell RNA-Seq library preparation protocols. Based on the performance, two RTases were further tested in high-throughput single-cell experiment.

**Results:** All RTases tested reverse transcribed low-concentration templates with high accuracy (*R^2^ > 0.9445*) but variable reproducibility (median CV_RT_ = 40.1 %). The most pronounced differences were found in the ability to capture rare transcripts (0 - 90% reaction positivity rate) as well as in the rate of RNA conversion to cDNA (7.3 - 124.5 % absolute yield). Finally, RTase performance and reproducibility across all tested parameters were compared using Z-scores and validity of obtained results was confirmed in a single-cell model experiment. The better performing RTase provided higher positive reaction rate and expression levels and improved resolution in clustering analysis.

**Conclusions:** We performed a comprehensive comparison of 11 RTases in low RNA concentration range and identified two best-performing enzymes (Maxima H-; SuperScript IV). We found that using better-performing enzyme (Maxima H-) over commonly-used below-average performer (SuperScript II) increases the sensitivity of single-cell experiment. Our results provide a reference for the improvement of current single-cell quantification protocols.

## INTRODUCTION

Single cell transcriptomics has emerged as a revolutionary technology transforming the biomedical research. From the technical perspective, minute amounts of RNA found in single cells have to be converted into cDNA, amplified and transformed into sequencing libraries. Reverse transcription (RT) is a critical step in this process, as any RNA molecules that fail to be initially captured are missing in the final data. Many factors can influence RT outcome, but reverse transcriptase (RTase) is arguably the most prominent. Wild-type RTases are not efficient and reliable enough for diagnostic and research applications. For *in vitro* purposes, many properties, such as thermostability, fidelity, processivity, substrate binding affinity, template switching activity and other properties are added or enhanced by engineering^1–4^. The alteration of these intrinsic properties is closely linked to reaction outcome. Currently, derivatives of Moloney Murine Leukemia Virus (MMLV) and Avian Myeloblastosis Virus (AMV) RTases are the most common, although other promising sources of RTases have been also identified^5^.

In addition to the selection of RTase also other factors must be considered when planning an experiment, such as priming strategy or template concentration^6,7^. Oligo(dT) primers target polyadenylated RNAs, while random sequence primers target all RNAs including the abundant rRNA fraction. For the ability to enrich for polyadenylated RNAs, oligo(dT)s have found the application in numerous RNA-Seq protocols^8–10^, whereas the mixture of random hexamers with oligo(dT)s are predominantly used in RT-qPCR experiments where it maximizes the reaction yield^11^. Gene-specific primers may be used as well to improve the specificity of the reaction. They are the most popular for targeted RT-qPCR applications mainly in diagnostics^6,7^ or for specific applications such as miRNA quantification^12–14^.

Experimental parameters influencing the conversion of RNA into cDNA were studied to a varying degree in past decades. The largest focus has been pointed on the choice of reverse transcriptase. Generally, MMLV-derived RTases have been found to deliver superior results^11,15–17^. Specifically, SuperScript II and SuperScript III (Thermo Fisher Scientific, USA) have been several times highlighted for their reproducibility and sensitivity^15,16,18,19^. However, not all RTases reported such constant quality. For example, performance of OmniScript and SensiScript (both Qiagen, Germany) or AMV derived RTases have been heavily influenced by experimental conditions, including the template input or by the laboratory conducting the experiment^15,17,18^. Notably, RTases with template-switching properties required for certain RNA-Seq protocols^4,20,21^ were recently compared by Bagnoli *et al.* scoring Maxima H- (Thermo Fisher Scientific, USA) and SmartScribe (Clontech, USA) as top performing candidates^9^. Several studies also focused on the role of other reaction components on the reaction outcome. The effectiveness of priming strategy was shown to be substantially gene-dependent^6,7,22^ but the discrepancies could be minimized by optimizing the primer concentration^19^. Gene-related efficiency of RT had been notified on multiple occasions^6,15,17,18,23^. Several reports suggested the addition of carrier molecules - tRNA^6^, polyethylene glycol^9^ or total extracted RNA^18^ to increase reaction yield.

Although efforts have been made to characterize the influences of aforementioned factors, little attention has been paid to characterize the performance of RT in low-RNA input applications, such as single-cell RNA-Seq. The exception is a study performed by Levesque-Sergerie *et al*. that partially mimicked single-cell conditions (detecting a single target in concentration of thousands copies per reaction)^18^. Unfortunately, the experimental extent as well as the number of tested RTases (five) were limited, therefore the study did not address many aspects of the RT for limiting template concentrations.

Here, we decided to fill the gap and benchmark a broad spectrum of currently available RTases in low-template applications. We systematically compared 11 RTases using equivalents of single-cell as well as 100-cell samples with two priming strategies popular for RT-qPCR and RNA-Seq applications. We characterized the RTases in terms of sensitivity, accuracy, yield and reproducibility. In the single-cell model study we demonstrated RTase influence on the experimental result quantified by differential positivity rates, higher expression levels and data clustering. Our results uncovered the substantial differences between individual RTases currently available and provide the data for an informed selection of the best suited RTase for the particular application in, but also outside the field of single-cell transcriptomics.

## METHODS

### Experimental design

The RTases were compared in two experiments: i) RTase benchmarking and ii) High-throughput validation (Figure 1). The goal of RTase benchmarking was to test the performance of 11 commercially available RTases (Table 1) in conditions mimicking single-cell and 100-cell samples. The follow up validation experiment aimed to illustrate the impact of RTase in gene expression profiling experiments using real single-cell samples. In both parts, two RT priming strategies were utilized: 1) equimolar mixture of 50 μM random hexamers with 50 μM oligo(dT)_15_ (recommended concentration in RT-qPCR protocols (*Supplementary protocols - Supp. data 1*)) and 2) 10μM oligo(dT)_15_ (recommended concentration in single-cell RNA-Seq protocols^9^).

**Figure 1.**
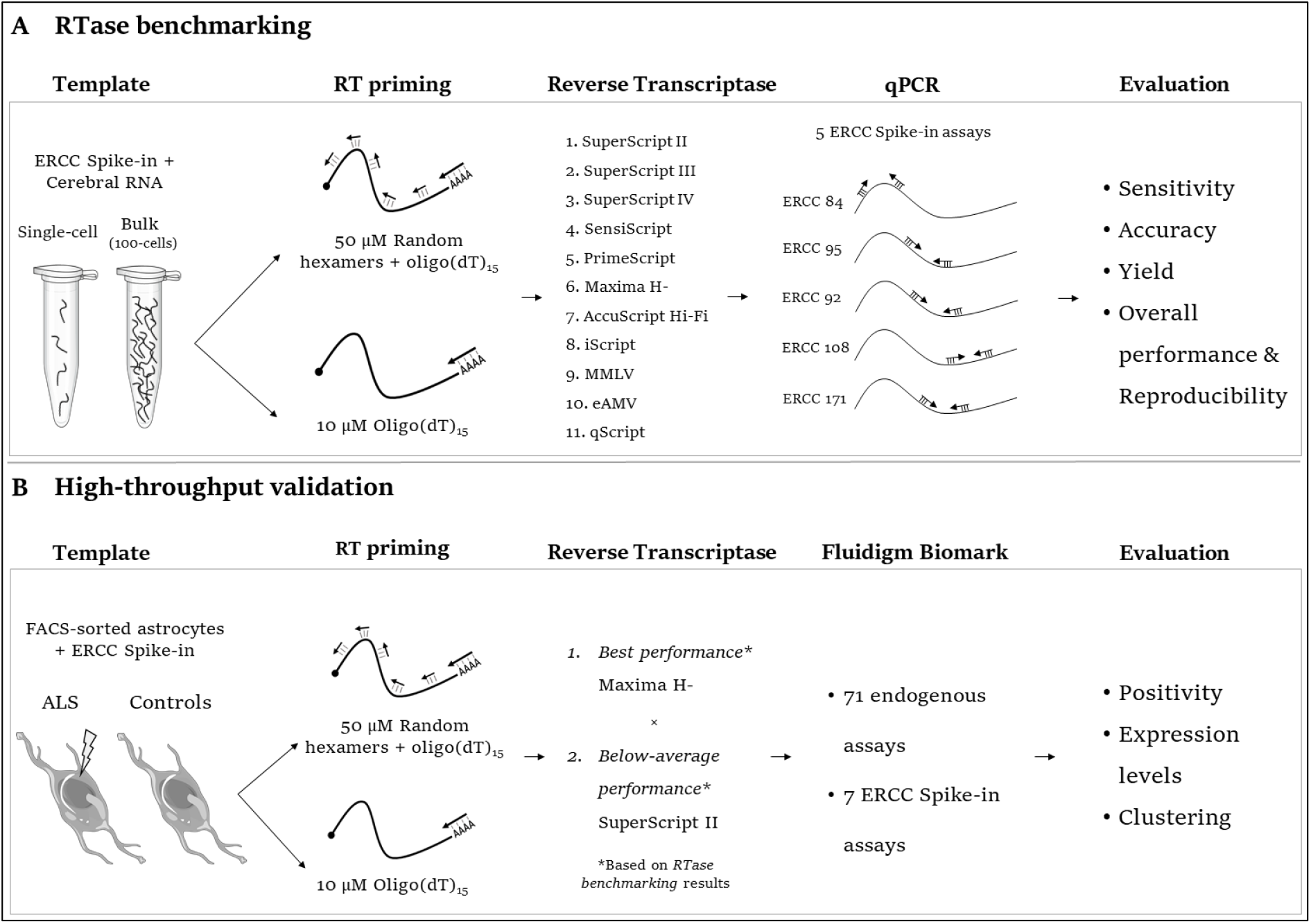
Experimental design.

**Table 1.** List of benchmarked RTases.

| Reverse Transcriptase | Manufacturer | Origin | RNase H activity | Reaction temperature (°C) | Cost per 20 μl reaction (\$) |
| --- | --- | --- | --- | --- | --- |
| SuperScript II | Thermo Fisher Scientific (USA) | MLLV | reduced | 42 | 7.5 |
| SuperScript III | Thermo Fisher Scientific (USA) | MLLV | reduced | 50 | 8.2 |
| SuperScript IV | Thermo Fisher Scientific (USA) | MLLV | reduced | 50 | 8.2 |
| Sensiscript | Qiagen (Germany) | unspecified | present | 37 | 6.0 |
| PrimeScript | Takara Bio Inc. (Japan) | MLLV | none | 42 | 6.5 |
| Maxima H- | Thermo Fisher Scientific (USA) | MLLV | none | 50 | 4.0 |
| AccuScript Hi-Fi | Agilent (USA) | MLLV | none | 42 | 12.0 |
| iScript | Bio-Rad Laboratories (USA) | MLLV | present | 42 | 6.5 |
| MLLV | Promega (USA) | MLLV | present | 42 | 2.0 |
| eAMV | Merck KGaA (Germany) | AMV | unspecified | 42 | 8.0 |
| qScript | Quanta Biosciences (USA) | MLLV | present | 42 | 5.2 |

### RTase benchmarking

#### Template

External RNA Controls Consortium (ERCC) Spike-in (set 1) (Thermo Fisher Scientific, USA) was used as primary template^24^. ERCC Spike-in set consists of unlabeled polyadenylated transcripts of various lengths (250 to 2000 nucleotides) mimicking eukaryotic mRNAs. Due to its known copy numbers and availability of DNA standards for absolute quantification of cDNA molecules, it was possible to calculate RNA conversion rate (see *Yield calculation, Methods* and *ERCC Spike-in validation, Supp. data 2* for details). Stock ERCC Spike-in was 200,000× diluted in TE-buffer supplemented with linear polyacrylamide (TE-LPA) for single-cell conditions or 2,000× for bulk conditions and stored in aliquots (Table 2). For each experiment a fresh aliquot was mixed with either 30 pg mouse cerebral RNA (used as a background) mimicking a single-cell or 3 ng mimicking 100-cell bulk sample^25^ (details in *RNA material*, *Supp. data 1,2*). In total, the ERCC Spike-in accounted for ~13 % of mRNA and ~0.5 % of total RNA, respectively.

**Table 2.** Assay specifications. Limit of detection is the lowest concentration that gives rise to positive signal in 95 % of cases. Limit of quantification is defined as the lowest concentration producing a standard deviation (SD) < 0.5 among replicates. Amount of nucleic acid in bulk samples was 100 times higher.

| Name | GenBank accession | Length [bases] | Single-cells: transcript count per RT | Single-cells: cDNA count per qPCR | Limit of Detection [molecules per qPCR] | Limit of Quantification [molecules per qPCR] | Efficiency |
| --- | --- | --- | --- | --- | --- | --- | --- |
| ERCC-00084-01 | DQ883682 | 970 | 22.0 | 1.8 | 5.7 | 6.4 | 0.95 |
| ERCC-00095-01 | DQ516759 | 495 | 88.2 | 7.1 | 5.7 | 100 | 0.863 |
| ERCC-00092-01 | DQ459425 | 1110 | 176.4 | 14.1 | 5.7 | 40 | 0.973 |
| ERCC-00108-01 | DQ668365 | 997 | 705.5 | 56.4 | 5.7 | 16 | 0.966 |
| ERCC-00171-01 | DQ854994 | 481 | 2821.9 | 225.8 | 5.4 | 6.4 | 0.978 |

#### Reverse transcription

For each RTase, reaction conditions and thermal profile followed the recommended protocol issued by manufacturer. General reaction components including nuclease-free water (NFW), RT primers, dNTPs, dithiothreitol (DTT), and ribonuclease inhibitor (all Thermo Fisher Scientific, USA) were supplied from a single stock. Total RT reaction volume was 5 μl. Each experimental condition was run in 10 replicates unless stated otherwise.

An exemplary RT reaction consisted of: 2 μl of background RNA (3 ng for bulk and 30 pg for single-cell equivalents), 0.5 μl of ERCC Spike-in (2,000× or 200,000× diluted) and 2.5 μl of RT mastermix containing RT primers (50μM RT primers mixture or 10 μM oligo(dT)_15_), 10mM dNTPs, RTase specific buffer, 0.1M DTT - if requested, 10 U of RNaseOUT - if requested, NFW and RTase). All RTs were performed in Bio-Rad C1000 Thermal Cycler (Bio-Rad, USA). Prepared cDNA was 5× diluted in NFW and directly used in qPCR to avoid freeze-thawing cycles. RTases included in this study are listed in Table 1. Detailed protocols are listed in *Supplementary protocols* (*Supp. data 1*).

#### Quantitative PCR

qPCR measurements followed a thoroughly validated protocol. A reaction volume of 10 μl contained 2 μl of 5× diluted cDNA and 8 μl of qPCR mastermix, which consisted of 2.6 μl NFW, 5 μl TATAA SYBR Green mix (TATAA, Sweden) and 0.4 μl of 10 μM primers (Thermo Fisher Scientific, USA). Cycling protocol consisted of initial enzyme activation at 95°C (t = 3 min), followed by 45 cycles of denaturation at 95°C (*t* = 15 s), annealing at 60°C (*t* = 20 s) and elongation at 72°C (*t* = 20 s). Melt curve analysis was performed in the temperature interval of 65 to 95°C, using gradient of 0.5°C. Bio-Rad CFX 384 (Bio-Rad, USA) thermal cycler was used for all measurements. CFX Manager Software (Bio-Rad, USA) and Project R were used for data processing.

The initial comparison of 11 RTases was performed using five thoroughly validated qPCR assays (Table 2). Detailed information on assay optimization and validation is listed in *ERCC Spike-in assay validation* (*Supp. data 1,2*). For each assay, three performance metrics were determined: efficiency (*E*), limit of detection (LOD) and limit of quantification (LOQ) (*LOD – LOQ* tab, *Supp. data 2*). The assays were selected to target ERCC molecules present at different abundancies (22 - 2822 copies per RT reaction in single-cell set-up) mimicking genes expressed at different levels.

#### Yield calculation

The quantification of absolute yield lies in determining relation between RNA input and cDNA output reported by qPCR. Input RNA copy numbers were calculated from the ERCC concentration provided by the manufacturer. The cDNA concentration was determined from the measured Cq and a DNA standard curve covering the range from 2 × 10^5^ to 2 × 10^-1^ copies (*n* = 4 replicates) (*ERCC Spike-in assay validation, Supp. data 2*). DNA standards were prepared from PCR-enriched target sequences. The RT yield is estimated using Equation 1:

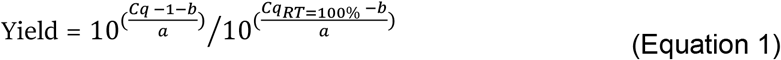

where *a* and *b* refer to slope and intercept of the particular ERCC Spike-in assay DNA standard curve, respectively and *Cq_RT=100%_* is the Cq expected for 100% RT efficiency. Adjusting Cq by −1 accounts for difference between single-stranded cDNA and double-stranded DNA standard used in standard curve construction.

### High-throughput validation on biological samples

#### Animals

All procedures involving the use of laboratory animals were performed in accordance with the European Community Council Directive of 24 November 1986 (86/609/EEC) and animal care guidelines approved by the Institute of Experimental Medicine, Academy of Sciences of the Czech Republic (Animal Care Committee decision on 17 April 2009; approval number 85/2009). Double transgenic mice bearing SOD1(G93A) and GFAP/EGFP alterations were used as a model organism. Mice with transgenic expression of mutant human superoxide dismutase SOD1(G93A) exhibit phenotype similar to amyotrophic lateral sclerosis (ALS) in humans, whereas GFAP/EGFP allows for visualization of astrocytes due to expression of EGFP protein under human glial fibrillary acidic protein (GFAP) promoter. EGFP-positive single-cells from dissected mouse tissue brain (3-months old) were collected using fluorescence-activated cell sorting GRISORBEON system 1 (FACS;0.9.21 BDInflux) into 5 μl lysis buffer (NFW + 1 mg/ml BSA). 96 single-cells were collected from both ALS and control mice. Details regarding the single-cell suspension preparation and cell sorting may be found elsewhere^26^.

#### Reverse transcription

Single-cells were reverse transcribed in a volume of 10 μl, consisting of: 5 μl of single-cell in lysis buffer, 0.5 μl of 200,000× diluted ERCC Spike-in and 4.5 μl of RT mastermix. RT mastermix preparation and thermal program followed RTase-specific protocol for Maxima H- and SuperScript II (*Supplementary protocols, Supp. data 1*). RT mastermix contained: RT primers (50μM RT primers mixture or 10μM oligo(dT)_15_), 10mM dNTPs, RTase specific buffer, 0.1M DTT (SuperScript II only), 20 U of RNaseOUT, NFW and RTase. Sample distribution followed orthogonal experimental design.

#### Samples – preamplification and quality control

All cDNA samples were preamplified in 40 μl total reaction volume comprising of 4 μl undiluted cDNA and 36 μl preAMP mastermix, which consisted of NFW, IQ Supermix buffer (Bio-Rad, USA) and 250 nM primer mix of 78 assays (Thermo Fisher Scientific, USA) (both endogenous and ERCC Spike-in assays). Detailed gene list is found in *Primer sequences* (*Supp. data 1*). Thermal protocol comprised of heating at 95°C (*t* = 3 min), followed by 18 cycles of template denaturation at 95°C (*t* = 20 s), annealing at 57°C (*t* = 4 min) and elongation at 72°C (*t* = 20 s). Template was immediately cooled on ice, 4× diluted in NFW and stored at −80°C.

4× diluted preAMP cDNA was used in qPCR quality control reactions. The expression of 4 genes were measured – *Gja1* (astrocyte marker), *Cspg4* (NG2 cells), *Vim* and *Slc1a3* (both commonly expressed in astrocytes). Only cells positive for *Gja1* and concurrently negative for *Cspg4* were used for qPCR analysis; remaining two genes were used as cell quality identifiers. preAMP reproducibility was also verified (*preAMP validation, Supp. data 1, 2*).

#### High-throughput qPCR

High-throughput measurements were conducted on a 96.96 Fluidigm BioMark platform (Fluidigm, USA). Protocol was as described in Rusnakova *et al*.^26^. The cycling program consisted of activation at 95°C (*t* = 3 min), followed by 40 cycles of denaturation at 95°C (*t* = 5 s), annealing at 60°C (*t* = 15 s) and elongation at 72°C (*t* = 20 s). After qPCR, melting curves were measured between 60°C and 95°C with 0.5°C increments. Results were post-processed based on melting curve inspection.

## RESULTS

### RTase benchmarking

In order to obtain a comprehensive view on the current state of commercially available RTases for single-cell applications, the enzymes were benchmarked on templates regularly used in single-cell RNA-Seq experiments (Figure 1 A). Single-cell or 100-cell bulk templates were primed with two priming strategies: RT primers mixture (common for RT-qPCR) or oligo(dT) priming (common for RNA-Seq). Performance of 11 RTases was assessed on 5 assays targeting ERCC Spike-in molecules of different abundancy, allowing to investigate the range from rare to highly-abundant transcripts (Table 2). Following the comparison on standardized samples, the performance of two RTases was further validated on a single-cell model experiment (*High-throughput validation* section) (Figure 1 B).

#### Sensitivity

To assess reaction sensitivity, the rate of positive reactions for low and medium abundant assays (*ERCC 84* and *95*) was determined. Reaction output was counted binary – positive or negative reaction, independent of signal strength. Two main parameters influenced reaction sensitivity – RT priming strategy and RTase itself.

Medium abundant assay *ERCC 95* (88 molecules per RT) reported little differences between priming strategies (Figure 2 A). For 9 out of 11 RTases recorded at least 80 % of the reactions positive with both priming strategies. Out of the two remaining enzymes - SuperScript II and eAMV, only the latter obtained less than 50 % of the reactions positive.

**Figure 2.**
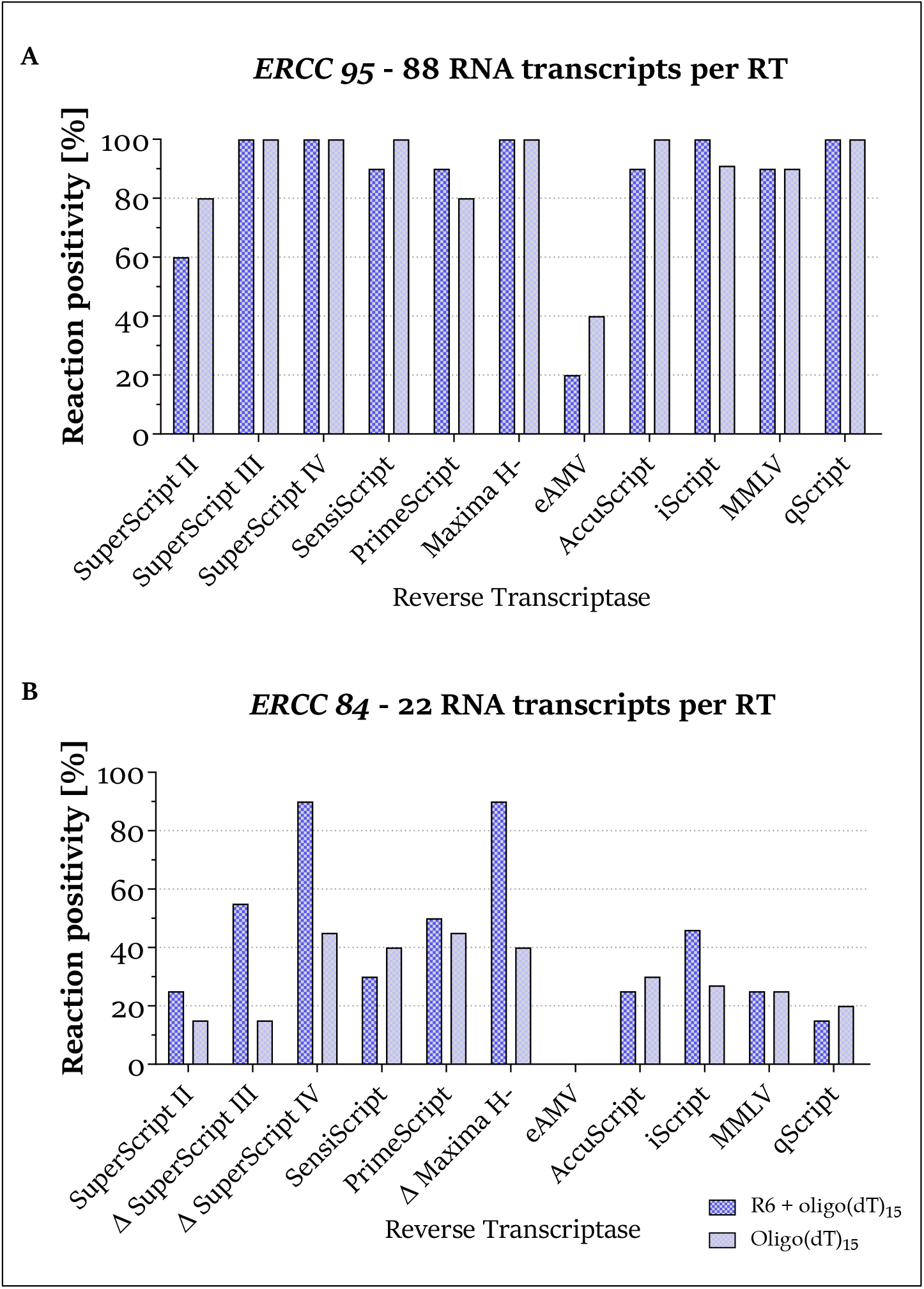
Sensitivity of the reaction is heavily influenced by the choice of RTase and priming strategy. RTases denoted with “Δ” doubled their rate of positive reactions with RT primers mixture. **A** Reaction positivity for medium abundant template (n = 10 reactions per RTase). **B** Reaction positivity for low abundant template (n = 20 reactions per RTase).

Four times less abundant template *ERCC 84* (22 molecules per RT) proved to be more challenging (Figure 2 B). The median positivity rate across all RTases was 30 %. Maxima H- and SuperScript IV stood out of the comparison as their sensitivity was strongly enhanced with RT primers mixture, reaching positivity rates of 90 %. With oligo(dT) priming their sensitivity dropped to ~45 %. Similarly, the rate of positive reactions with SuperScript III dropped by more than half when primed with oligo(dT). Notably, with SuperScript II, recommended in some RNA-Seq protocols^20,27^, only ~20 % of the reactions were positive. The only AMV-derived RTase in our comparison, eAMV, failed to show any positive reactions.

#### Accuracy

Reliable and unbiased RT outcome is critically important for every study. Inconsistent RT efficiency at any template concentration could potentially lead to false conclusions. To assess the accuracy of the RTases over a wide concentration range, we merged single-cell and 100-cell measurements and plotted RNA-molecule concentration (22 to 282,000 specific ERCC Spike-in copies per RT reaction) versus cDNA concentrations estimated by qPCR. Absolute cDNA concentrations were estimated using Equation 1 (see *Methods*), which accounts for assay-specific differences in efficiency. Coefficients of Determination (*R^2^*) served as metric of accuracy.

All studied RTases reported accurate performance with both priming strategies. *R^2^* for the RT primers mixture ranged from 0.9463 to 0.9896, and with oligo(dT)_15_ priming the values ranged from 0.9445 to 0.9825. Reproducibility varied considerably between single-cell and bulk templates, however the expected performance remained linear across tested RNA input range (Two representative RTases shown in Figure 3; remaining results are listed in *Accuracy – linearity* tab, *Supp. data 1*). When looking at the reproducibility of RT replicates for each RTase separately, the major effect of template abundancy was identified. Among tested RTases, the number of cDNA copies per single-cell reaction varied in median by 40.06 % (coefficient of variation - CV_RT_), while for bulk samples median variation was substantially lower at CV_RT_ = 10.27 %. The most reproducible RTases in single-conditions were Maxima H- and SuperScript IV with median CV_RT_ of 29.15 % and 30.25 %, respectively. The least reproducible RT reactions were observed for SuperScript II (CV_RT_ = 53.65 %) (*Reproducibility* tab, *Supp. data 1*). In conclusion, although reproducibility of RT decreases with template concentration, RT is overall substantially accurate even at single-cell levels regardless of used enzyme or priming strategy.

**Figure 3.**
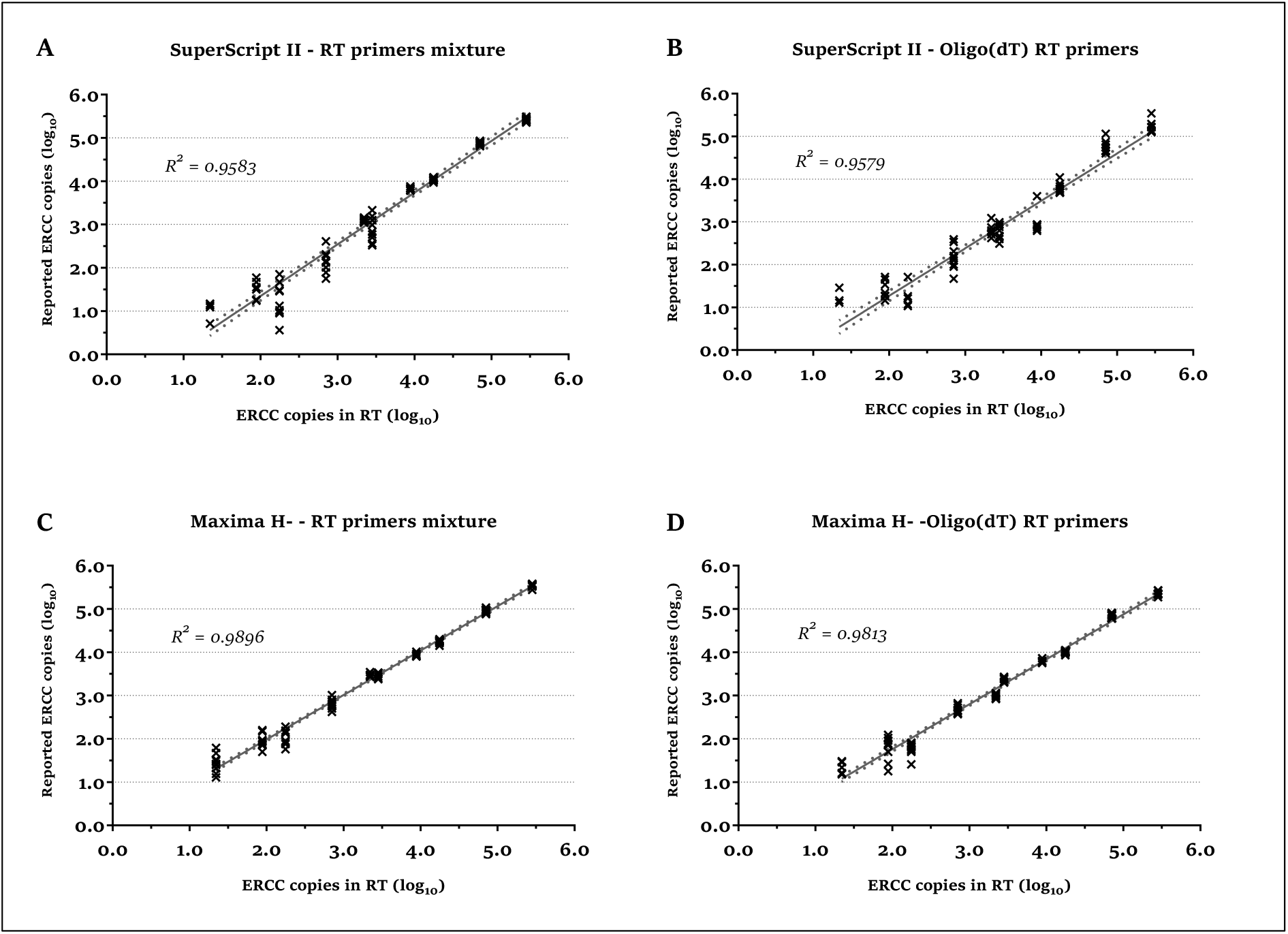
RT performs with substantial accuracy even at single-cell conditions. Linear regression plots reporting log_10_(captured ERCC copies) versus log_10_(input ERCC copies) for SuperScript II (top) and Maxima H- (bottom) using RT primers mixture (left) and oligo(dT) primers (right). Graphs display the best linear fit and its 95 % confidence intervals. R^2^ reflects the quality of fitted regression line. Reproducibility decreases at lower template concentrations.

#### Yield

Reaction yield is one of the most critical performance metrics. Ideally the reaction would approach 100 % value, however, previous studies have shown that the actual rate varies substantially^15,18,19^. The use of ERCC Spike-ins and DNA standards enabled us to quantify yields based on known target copy numbers per reaction (Equation 1, see *Methods*).

For single-cell template, significant differences in yield were observed between RTases and priming strategies (*two-way ANOVA, p_RTase_* < *0.001, p_priming_* = *0.005 and p_interaction_* < *0.001*). Reactions primed with RT primers mixture (Figure 4 A) were best processed by SuperScript IV, Maxima H- and SuperScript III, as they reported average yields of 125 %, 102 % and 88 %, respectively. Theoretically, RTases with the strand displacement capability could produce multiple cDNA copies from a single RNA transcript, thus reaching yields over 100 %. The lowest yields with RT primers mixture were 24 % and 7 % for SuperScript II and eAMV, respectively. With polyA priming, there was no RTase significantly outperforming all others (Figure 4 B); SuperScript IV and Maxima H- showed the highest (71 % and 66 %) and eAMV the lowest yield (14 %). To remove the influence of priming strategy and identify the differences between RTases, *one-way ANOVA* was used separately for each priming strategy. The effect of RTase was more pronounced for reactions with RT primers mixture (*one-way ANOVA*, explained variation – *R^2^* = 0.8212) compared to polyA-targeted priming (*one-way ANOVA, R^2^* = 0.6657). Significant mean differences based on the Tukey *post-hoc multiple comparison test* (Tukey HSD) are indicated with letters in Figure 4.

**Figure 4.**
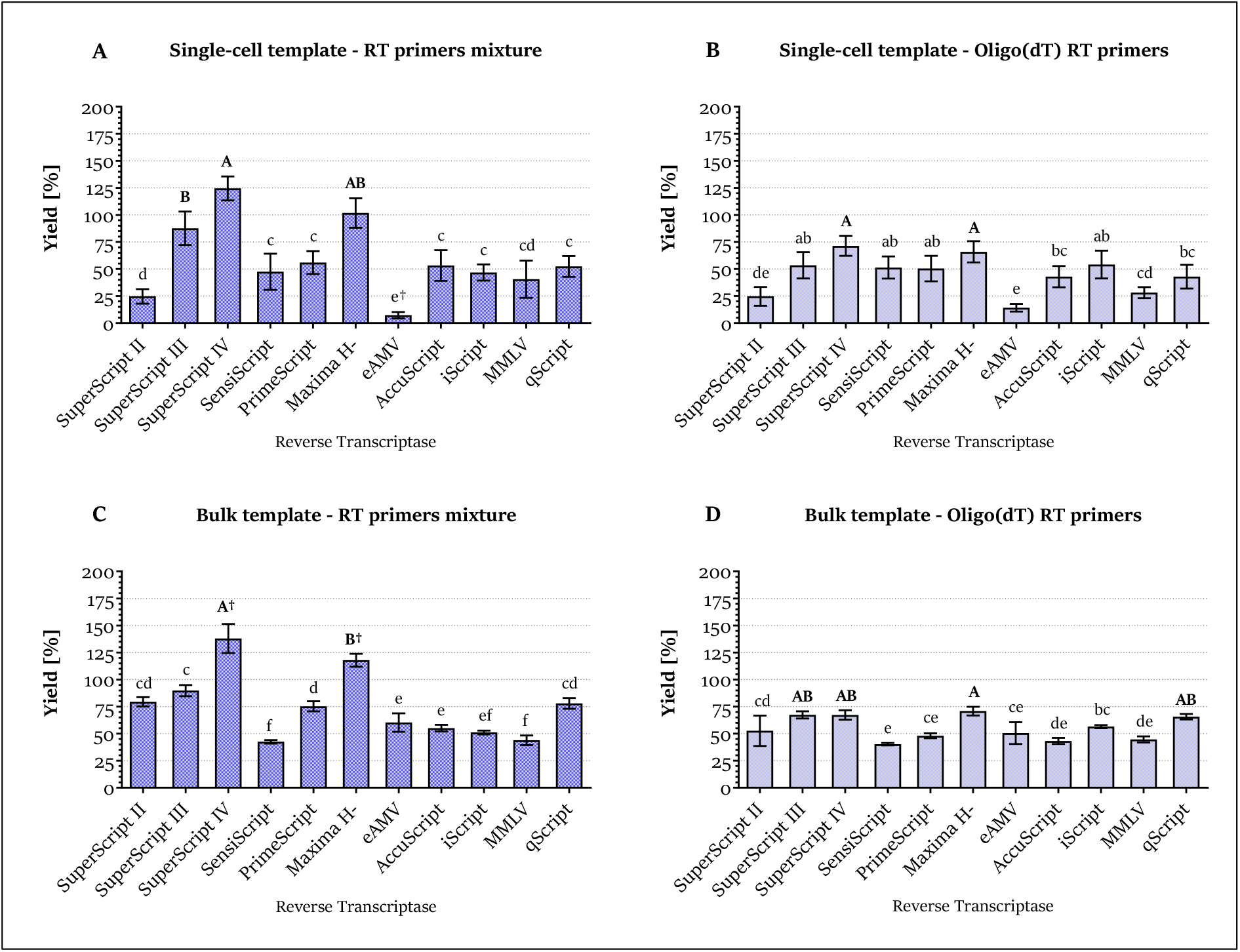
Reaction yield significantly depends on the choice of RTase and priming strategy. Bar plots show average yield per n = 10 RT replicates (1 RT replicate = average yield across 5 assays) with 95 % CI. Letters indicate significant mean differences between RTases (p < 0.05, Tukey HSD post hoc comparison), with the highest yield in bold. “†” marks unique significantly different mean value.

For bulk samples, SuperScript IV and Maxima H- showed the highest yields of 138 % and 118 % with RT primers mixture, respectively (Figure 4 C). On the other side, the lowest yields with RT primers mixture were reported with MMLV and SensiScript enzymes (44 % and 43 %), respectively. With oligo(dT) primers, Maxima H-, SuperScript IV, SuperScript III and qScript were the best scoring RTases yielding 71 %, 67 %, 67 % and 66 %, respectively (Figure 4 D). SensiScript recorded the lowest yield of 40 %. Differences between RTases were more prominent with RT primers mixture (*one-way ANOVA, p < 0.001, R^2^* = 0.9319) than with oligo(dT)s (*one-way ANOVA, p < 0.001, R^2^ = 0.674*).

Overall, the highest cDNA synthesis yields were obtained with SuperScript IV and Maxima H-. In the opposite spectrum, eAMV was found to be the least yielding RTase. The application of RT primers mixture resulted in the total yield over 100 % for some RTases, while with oligo(dT) the maximal value was capped around 75 %.

#### Performance reproducibility

To compare the overall performance of RTases relative to each other and independently of transcript abundancy, we applied Z-score scaling^28^. Cq-derived Z-scores in the form of boxplots (Figure 5) indicate the general RTase performance across all tested parameters (higher Z-score indicates better performance), as well as the reproducibility of such performance (spread of values). To highlight the difference in the reproducibility, Z-scores across priming strategies were combined.

**Figure 5.**
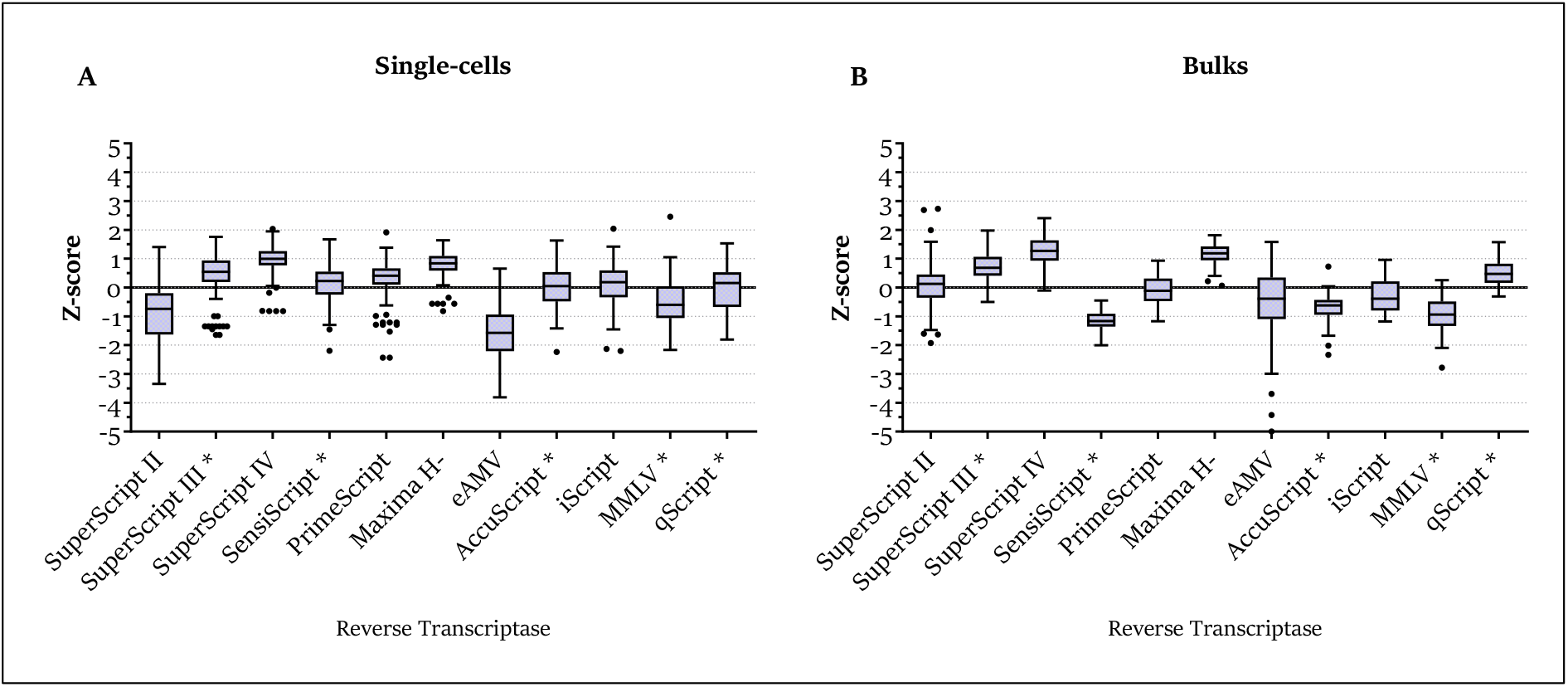
Best performing RTases retain consistently better performance even in single-cell conditions. Z-scores inform about the reproducibility and overall relative performance of RTases in single-cell **(A)** and 100-cell reactions **(B)**. Unit of 1 Z-score equals to difference of 1 SD from the assay’s average Cq for given template and given priming strategy. For enzymes denoted with “*”, performance with single-cell template was significantly less reproducible than with bulk samples. Performance of the best RTases (scoring the highest Z-scores) was found to be consistent, seen as small interquartile range.

Maxima H- and SuperScript IV were the best performers among the studied RTases. A median Z-score of 0.99 (interquartile range: 0.78 – 1.25) and 0.84 (0.59 – 1.09) was obtained for single-cell measurements for SuperScript IV and Maxima H-, respectively. For bulk measurements, SuperScript IV and Maxima H- reported to have median Z-score of 1.27 (0.94 – 1.62) and 1.18 (0.95 – 1.43), respectively. Among the other RTases, eAMV and SuperScript II were the least suitable for single-cell conditions, scoring −1.58 (−2.21 − −1.02) and −0.74 (−1.59 − −0.23) respectively.

The performance reproducibility varied significantly among tested RTases as well (*classical Levene’s test, p < 0.001* for both templates). The least variable performance, measured as Z-score interquartile range, was recorded for SuperScript IV, Maxima H- and PrimeScript in single-cells (Figure 5 A). In bulk samples, SensiScript, Maxima H- and AccuScript were the most robust performers (Figure 5 B). Template abundancy had a significant impact on the performance reproducibility of five RTases (indicated with “*” in Figure 5; *classical Levene’s test, p < 0.05, adjusted with Bonferroni’s correction*).

In summary, RTases significantly varied in their performance and reproducibility. Among tested enzymes, Maxima H- and SuperScript IV were found to be superior to their counterparts in both tested parameters and may be recommended for low RNA input applications in a wide range of reaction conditions.

### High-throughput validation

The initial comparison of the 11 RTases allowed us to classify them based on their performance. However, the comparison was based on artificial templates (ERCC Spike-in) and the limited number of assays. Therefore, we evaluated two RTases, Maxima H- and SuperScript II, in a routine high-throughput single-cell RT-qPCR profiling experiment based on 78 assays (Figure 1 B). The two RTases were selected based on their performance (best-performer vs below-average performer) and also based on their routine application in single-cell RNA-Seq protocols^4,9,10,20,27^. In consistency with the first part of this study, two priming strategies (RT primers mixture and oligo(dT)) were compared.

FASC-sorted astrocytes from healthy and ALS mouse brains were used as single-cell samples. 15 wild-type and 5 ALS cells were analyzed with each RTase and priming strategy (Figure 1). To minimize the biological variability, the quality of cells was pretested and only cells passing the criteria were used for follow-up analysis (see *Materials and methods*). Additional quality filter was performed after data acquisition (negative assays were discarded, followed by melting curve analysis of assay specificity and control for astrocytic markers - *Gja1, Slc1a3* and *Aqp4*). After data pre-processing, we evaluated the RTases and priming strategies on positive call rate, expression levels and cluster separation.

#### Positive call rate and expression levels

The number of positive reactions with each RTase and priming strategy was evaluated. Maxima H- showed overall more positive reactions than SuperScript II (Figure 6 A, C). When using RT primers mixture, Maxima H- yielded more positive calls for 55 % of the assays (positivity rate increased on average by 12 % per assay); equal positive rate was found for 25 % of the assays and SuperScript II showed more positive calls for 20 % of the assays (Figure 6 A). Using oligo(dT)s the difference in RTase performance was magnified. Maxima H- gave more positive calls for 74 % of the assays (on average 15 % more positive reactions), 17 % of the assays had equal positive call rate and SuperScript II had more positive reactions only for 9 % of the assays (Figure 6 C).

**Figure 6.**
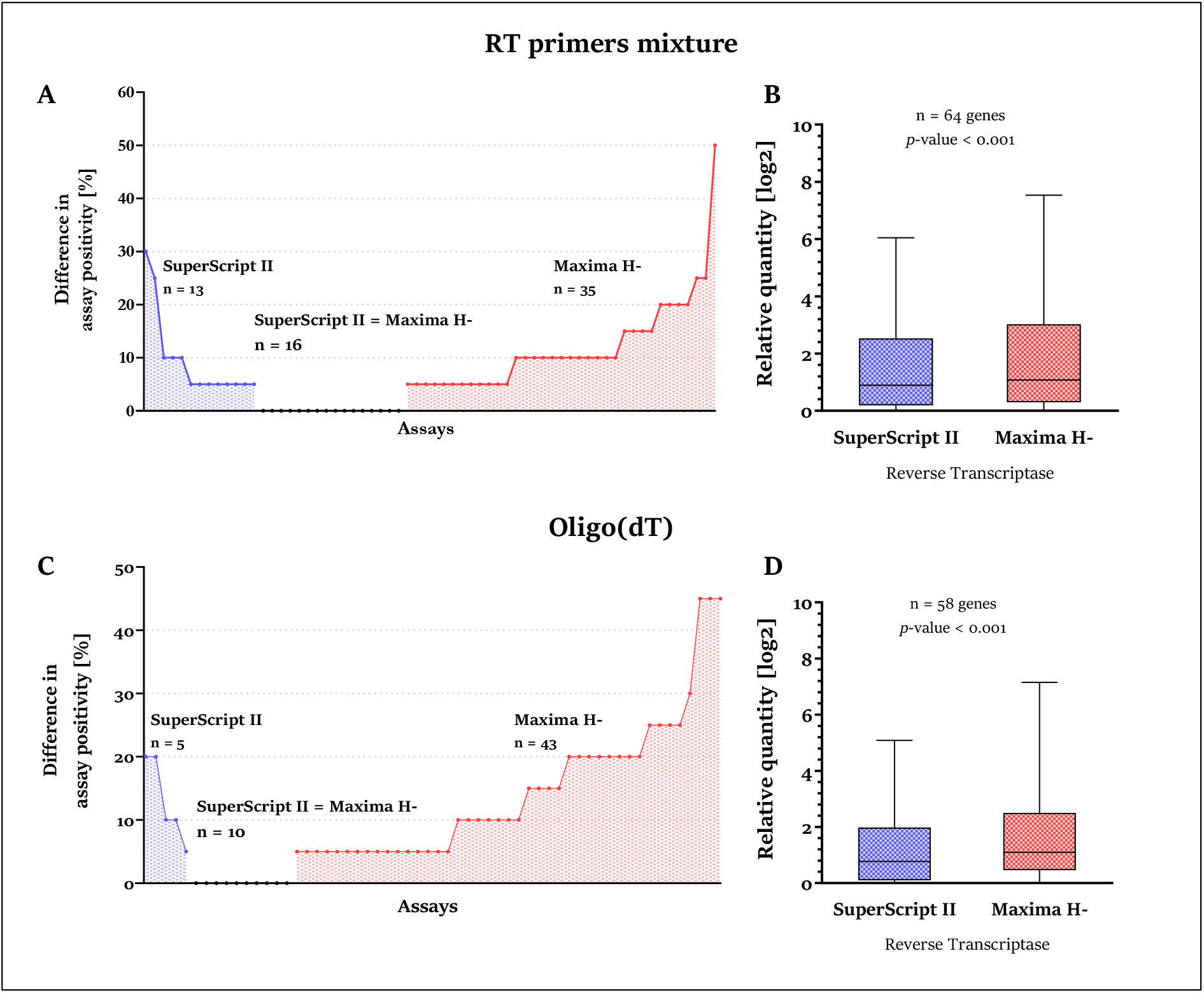
Maxima H- delivers more positive reactions and stronger detection signal. The RTase influences the amount of information obtained in high-throughput single-cell measurements (**A**,**C** – RT-qPCR reaction positive call rate; **B**,**D** – Expression levels; Tukey boxplots).

RTase and priming strategy have impact not only on the reaction positivity but also on the measured quantity as reflected by the Cq values. Consequently, RTase delivering stronger signal is more favorable in both qualitative and quantitative applications. The expression data was processed as relative quantities (RQs) calculating the level for each assay relative to the least abundant reaction within the assay, for each RT priming strategy. Significantly higher expression levels were obtained with Maxima H- with both priming strategies (Wilcoxon signed rank test, *p < 0.001*) (Figure 6 B,D).

#### Clustering

High-throughput gene expression experiments are typically analyzed using multivariate statistics^29,30^. To test if the RTase also impacts the cluster separation, we applied the common multivariate tool Principal Component Analysis (PCA). PCA was performed on Cq values that were firstly transformed to RQs relative to the least expressed sample and then converted to log_2_ scale. Redundancy analysis (RDA) identified significant importance of the RTase for the clustering for both priming methods, although values of explained variability (*R^2^*) were low (*RDA - covariate: enzyme; p = 0.036, R^2^ = 0.0386* for RT primers mixture and *p < 0.001, R^2^ = 0.0825* for oligo(dT)s). As expected, larger proportion of the total variance in the data was due to cell condition (control vs ALS). RT primers mixture recorded *R^2^* = 0.241 and oligo(dT)s *R^2^* = 0.28 (*RDA – covariate: cell treatment*). The impact of the RTase may be also quantified by the Euclidean distances between the two clusters’ centroids. When using the RT primers mixture, the cluster distance was ~20 % larger when using Maxima H- compared to SuperScript II (16.69 to 13.64 scores) (Figure 7 A). Using oligo(dT) priming, Maxima H- separated clusters ~40 % further than SuperScript II (18.23 and 12.97 scores, respectively) (Figure 7 B).

**Figure 7.**
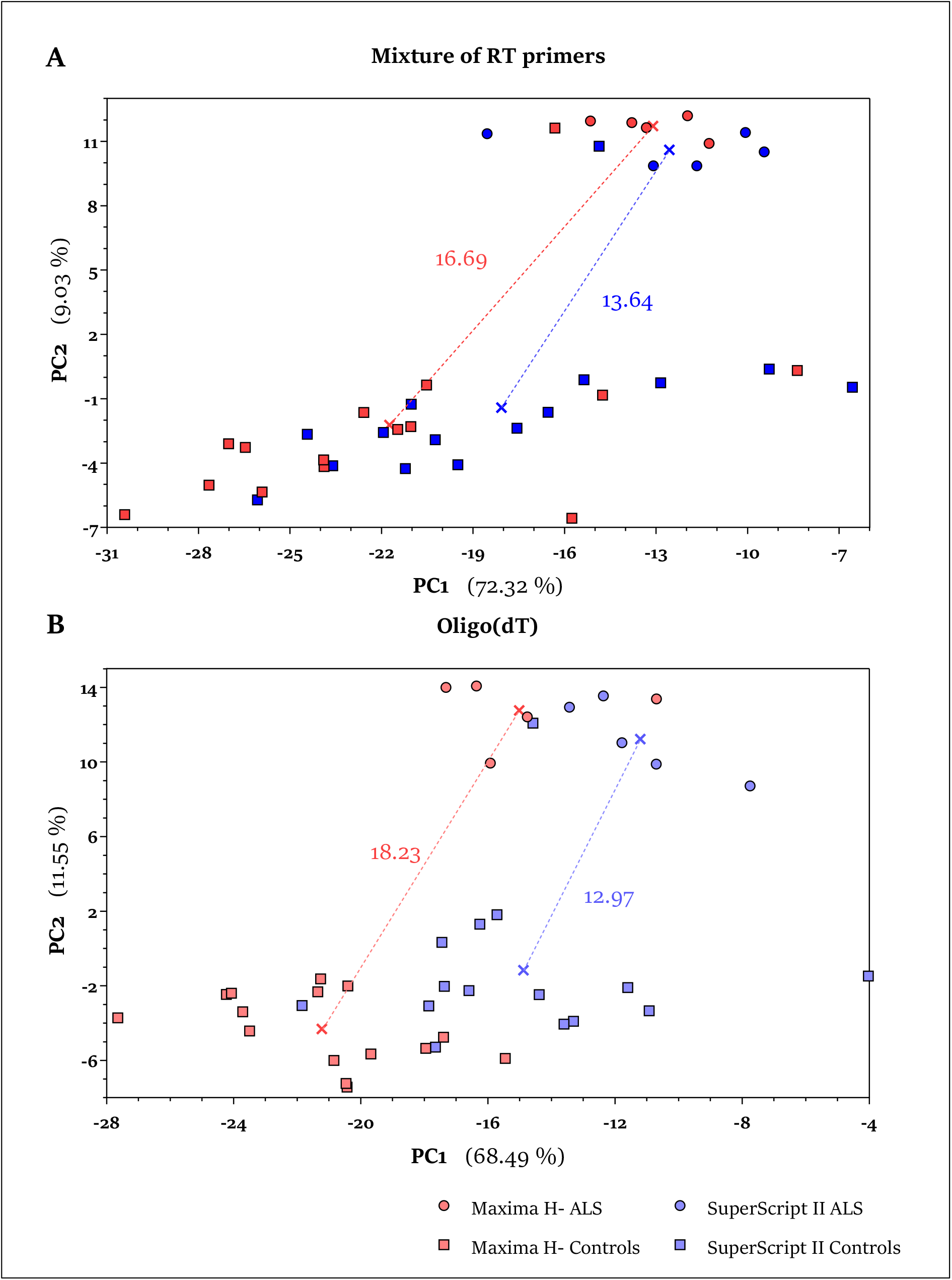
Better performing RTase separates biologically different clusters further apart. Using Maxima H-clusters of control and diseased cells separate better compared to SuperScript II. Accounted variance by the PCs is indicated in brackets. Cluster centroid is shown as color-adjusted cross.

In summary, the high-throughput experiment demonstrated the impact of RTase on the outcome of a typical single cell study as well as validated our previous data achieved using synthetic RNA template. The better performing enzyme achieved higher positive rate and signals as well as contributed to better separation of two populations of cells.

## DISCUSSION

In this study, we systematically evaluated the performance of 11 commercial RTases with focus on RNA-based single-cell quantitative protocols. Key findings include that: (i) accuracy of RT is generally high over wide range of template concentrations regardless of RTase and priming strategy; (ii) the sensitivity of majority of tested RTases is sufficient and comparable for more-abundant templates, but only few maintain sensitivity with low-abundant template; (iii) the high sensitivity of these RTases is accompanied with high reproducibility and therefore are preferable for single-cell applications; (iv) RTase performance is dependent on priming strategy; (v) usage of better RTase significantly improves the gene detection rate from single cells and improves definition of biologically distinct single-cell populations.

For single-cell expression profiling, the ability of RTase to capture transcripts present in low concentrations is of prime importance. Using a set of spike-in molecules enabled us to test this parameter over a wide range of concentrations (22 to 2822 molecules per RT reaction). With exception of eAMV, the reaction positivity rate of the RTases does not change considerably over the medium-to-high concentration range (>88 transcripts per reaction; Figure 2 A), but differs substantially for low-abundant transcripts (Figure 2 B). Considering that genes are expressed on median between ~3 and ~100 copies per cell^31^ (~100 copies for protein-coding, ~10 for splicing regulators and only ~3 for transcription factors), our results show that the choice of RTase represents a strong variable in the sensitivity of single-cell quantitative experiments. As the detection of low-abundant transcripts may be affected by sampling noise, the increased number of replicates we used for lowest-abundant transcript (*n* = 20) should minimize this factor and capture the trends in relative RTase performance. Importantly, the influence of sampling noise is insignificant over 35 template copies^32^, which highlights the poor performance of the AMV-based enzyme (Figure 2 A). Poor AMV performance has been noted before^11,15–17^ and has been attributed to its dimeric structure^33^.

Comparing priming strategies, we observed increased positivity rate (Figure 2 B) and increased yield (Figure 4 A,C) when using RT primers mix compared to oligo(dT) only. However, two confounding factors may contribute to this conclusion. First, polyA-based priming of ERCC Spike-in is known to underestimate real oligo(dT) priming efficiency^8^ as the ERCC Spike-ins have unnaturally short polyA tails (20-30 bp)^24^. Secondly, different concentration of RT primers mixture and oligo(dT) primers only was used throughout our study (to mimic their typical usage in RT-qPCR and RNA-Seq studies) which may contribute to the lower positivity rate as well^19,34^. Therefore, although different outcomes using two priming strategies are apparent, we caution against the simplified interpretation that the RT primers mix always outperforms oligo(dT) primers^6,35^.

The ability of RTases to reverse transcribe mRNA templates with the same efficiency irrespective of template concentration was tested on a range from 22 to 282,200 ERCC Spike-in molecules per RT. All enzymes retained linear performance across tested range (Figure 3, *Accuracy – linearity, Supp. data 1* tab), demonstrating that RTase can reliably reflect RNA content for templates of varying abundancy^8^, with particular enzyme-specific efficiency (see *Yield*, Figure 4). The variability of RT yield however increases with decreasing template concentration (*Reproducibility* tab, *Supp. data 1*)^11^. Although all RTases performed with substantial accuracy, better performing RTases showed better efficiency, were more reproducible and sensitive at single-cell level, which makes them more suitable candidates for single-cell applications. Some early reports proposed that RT efficiency is gene- or template concentration-dependent^6,15,34,35^. Our findings are however in accordance with more recent reports^8,11^ that suggest RT is comparably efficient for different assays and template concentration.

The RT yield showed large variation across tested enzymes and was priming strategy-dependent (Figure 4). More pronounced differences between yields of various enzymes with RT primers mixture suggest that especially SuperScript IV and Maxima H- can markedly benefit from increased number of priming locations^35^. Multiple priming locations in combination with strand displacement activity, lack of RNase H function, high processivity and strong template binding affinity enable synthesis of more than just one cDNA copy from one RNA transcript, thus yields > 100 %.

Differences in RT efficiency between single-cell and bulk templates were minor and relatively consistent with exception of SuperScript II and eAMV enzymes (Figure 4). The worse performance observed with SuperScript II and eAMV on the single-cell template could be expected, as the RNA amount was below the recommended template range by the manufacturers. Considering this, it is surprising that several popular single-cell RNA-Seq protocols (CEL-Seq2^27^, Smart-Seq2^20^) are based on SuperScript II, suggesting there is a potential for their improvement. Apart the discussed high and low efficient RTases, the cDNA synthesis yield was typically in the range 50 – 80 %, which is in line with earlier studies^8,15–17,23,34^. Only contradictory result is reported by Levesque-Sergerie *et al*. on SuperScript II and III RTases^18^, which may be attributed to imprecise preparation of qPCR standard curves as discussed by Miranda & Steward^19^.

Our study also reveals the relationship between performance of particular RTases and their intrinsic biochemical properties (Table 1, Figure 5). The best-performing RTases, SuperScript IV and Maxima H-, were thermostable, allowing to utilize higher reaction temperature in the protocols. Previous reports have suggested that destabilization of secondary RNA structures at elevated temperature leads to more frequent primer hybridization and stable reverse transcription^6,36^. Our results support these hypotheses. Interestingly, the majority of RTases employed protocols with the pre-incubation step aiming to increase the efficiency of primer binding. The exception in our selection was iScript and SensiScript. Both enzymes achieved relatively lower performance demonstrating the importance of the pre-incubation step. However, if loss of sensitivity is not an issue, simplified pipetting protocol, reduced possibility for contamination and errors may be advantageous factors. RNase H activity was not observed to have any profound effect on the reaction outcome, as previously reported^11,17^. The important factor for selection of RTase is also its price. This is relevant especially for single cell RNA-Seq studies, where thousands of cells are typically analyzed and price for RT is an important part of the budget. From this perspective, Maxima H- may be a recommendable choice for many researchers for its high performance and low price (second lowest in our comparison, Table 1). Notably, Maxima H- possesses also the terminal transferase activity utilized in some RNA-Seq protocols^8–10^ (mcSCRB-Seq, STRT-Seq, SMARTer, Smart-Seq2).

Typically, high-throughput experimental results require reduction of their multidimensional composition. PCA, the commonly used method, reduces dimensionality by searching for the largest portions of variation in the data set. Observed variation, however, arises not only from the biology of the experiment but also from technical factors of the measurement, including RT^30^. In our model experiment, we hypothesized that higher sensitivity will enhance separation of biologically distinct single-cell clusters by reducing the frequency of missing values and sampling noise^32^. Indeed, the number of positive reactions per assay as well as the expression levels were increased using Maxima H- compared to SuperScript II regardless of the priming strategy (Figure 6). The improvement is also reflected by increase in the cluster distances, as hypothesized (Figure 7). To our knowledge, this is the first study that demonstrated the direct effect of RT on separation of distinct groups of single cells in multidimensional expression analysis.

The aim of our study was to evaluate the current state of commercially available RTases for the growing field of single cell transcriptomics. We showed substantial differences in the performance of RTases highlighting the importance of their selection. For the first time, we also demonstrated the impact of RTase on the outcome of a typical single cell study. We believe that this study will initiate follow-up efforts to characterize other aspects of the RT reaction and will lead to the improvements of existing workflows.

## Acknowledgments

We would like to acknowledge the funding sources for this work: P303-19-02046S, CZ.1.05/1.1.00/02.0109 and RVO 86652036.

