## Supplementary data 2 for "Performance comparison of reverse transcriptases for single-cell studies"

### Supplementary methods

#### *RNA material*

Mouse tissue samples from cerebellum were dissected, placed into TRI Reagent (Sigma-Aldrich) and immediately frozen on dry ice, as previously described in Rusnakova *et al*[1]. Samples were stored at -80°C. Before use, samples were thawed, homogenized using the TissueLyser (Qiagen) and total RNA was extracted with TRI Reagent (Sigma-Aldrich) according to the manufacturer′s protocol. Samples did not undergo DNAse treatment. RNA quantity and purity were assessed using the NanoDrop 2000 spectrophotometer (Thermo Fisher) and RNA integrity was assessed using the Fragment Analyzer - DNF 489 Standard Sensitivity RNA Analysis Kit (Advanced Analytical Technologies, USA) (*RNA material* tab, *Supp. data 1*). To prepare aliquots, extracted RNA was diluted in nuclease-free water (NFW) to 1.5 ng/µl and then stored at -80°C. All procedures involving the use of laboratory animals were performed in accordance with the European Community Council Directive of 24 November 1986 (86/609/EEC) and animal care guidelines approved by the Institute of Experimental Medicine, Academy of Sciences of the Czech Republic (Animal Care Committee decision on 17 April 2009; approval number 85/2009). Identical set of RNA aliquots was used for both assay validation and RTase benchmarking. Capillary electrophoresis results (Fragment analyzer) are listed in *RNA material* tab (*Supp. data 1*).

#### *ERCC Spike-in assay validation*

qPCR primer sequences were designed using NCBI Primer Blast tool. NetPrimer (Premier Biosoft, USA) and OligoAnalyzer Tool (Integrated DNA technologies, USA) were used to check *in-silico* for primer-dimer formation. Validation of assays was performed in 2 steps: 1) General testing of primer specifity and 2) Standard curves/efficiency.

cDNA necessary for validation of assays was made in 10 µl reactions using SuperScript IV protocol (*Supplementary protocols* tab, *Supp. data 1*). Each assay was validated on samples: 1) four replicates containing 4 µl of cerebral RNA (1.5 ng/µl) spiked-in with 1 µl 100× diluted ERCC Spike-in, 2) one replicate of 2 µl mouse gDNA (0.5 ng/µl) and 3) three replicates of 2 µl nuclease-free water (NFW). RT was primed using 50 µM equimolar mixture of random hexamers with oligo$(dT)_{15}$. After RT, cDNA was 10× diluted. Remaining reagents were used according to *Supplementary protocols* tab (*Supp. data 1*). Amplification and melting curves are posted in *ERCC Spike-in assay validation* tab (*Supp. data 1*). Size and purity of PCR amplicons was verified by capillary electrophoresis (Fragment Analyzer – dsDNA 905 Reagent Kit, 1 bp – 500 bp). Results of PCR product size control are listed in *ERCC Spike-in assay validation* tab (*Supp. data 1*).

Diluted PCR products from the first round of primer testing (pool of two cDNA samples) were consequently used as a template for standard curves. The copy number of standard stock solutions was calculated using the concentration determined by Qubit 2.0. Qubit's 500 ng/ml standard was diluted into dilution series of 500 – 250 – 125 – 62.5 – 31.25 – 0 ng/ml. Linear regression (template abundancy ~ fluorescence signal) determined slope and intercept, which was used for calculation of concentration of unknown samples, i.e. PCR products. Based on known concentration of PCR molecules and theoretical weight of one double stranded ERCC Spike-in PCR product molecule, we calculated the number of molecules per 1 µl of sample. This information was used for dilution of the PCR products to required concentration, which was necessary for standard curve construction. Standard curves were performed in replicates of four. PCR products were diluted in TE-buffer supplemented with linear polyacrylamide (TE-LPA) to cover range from $2 \times 10^5$ to $2 \times 10^{-1}$ copy numbers per qPCR reaction. Standard curves are posted in *ERCC Spike-in assay validation* tab (*Supp. data 1*).

*Limit of Detection and Limit of Quantification (LoD and LoQ)*

LoD and LoQ were calculated using the same PCR products that were used for standard curve construction. For each assay, separate dilution series was prepared: 100 – 40 – 16 – 6.4 – 2.6 – 1 – 0.4 – 0.16 copies per qPCR reaction. Measurements were collected in qPCR replicates of six. Limit of Detection is defined as template concentration with 95 % confidence to be detected in all undergoing qPCR reactions. Limit of Quantification was defined as template concentration with SD(Cq) < 0.5 for all reactions. LoD and LoQ results are listed in *LoD – LoQ* tab (*Supp. data 1*).

*preAMP validation*

cDNA used in preAMP validation was made in 20 µl RT reaction using SuperScript II RTase protocol. In three RT replicates, the reactions contained 5 µl of 50 ng/µl cerebral RNA and 0.25 µl of 1000× diluted ERCC Spike-in. RT- reaction was also prepared. cDNA was immediately 4× diluted in NFW. Thermal protocol is described in *Supplementary protocols* tab (*Supp. data 1*).

preAMP reactions were prepared in 40 µl volume. Reactions containing 4 µl of 4× cDNA were prepared in duplicates. List of primers for preAMP mix is to be found in *Primer sequences* tab (*Supp. data 1*). Thermal protocol and reagents are described in *Supplementary protocols* tab (*Supp. data 1*). Preamplified cDNA was immediately 50× diluted in NFW and stored at -80°C.

Reproducibility of preAMP was validated by measuring the Cq difference ($\Delta AMP$) between preAMP-ed and non-preAMP-ed cDNAs. Both preAMP and non-preAMP cDNAs were used in 50× dilution as qPCR templates. Samples were measured in qPCR duplicates. qPCR followed protocol described in *Supplementary protocols* tab (*Supp. data 1*). Measured assays included high, medium and low abundant targets and both endogenous and ERCC Spike-in assays. Results for seven targets are listed in *preAMP validation* tab (*Supp. data 1*).

*Data analysis*

CFX Manager Software (Bio-Rad, USA) and Project R were used for data processing. All qPCR measurements were brought to uniform threshold value. Missing values were replaced with maximum Cq + 2 per each combination of assay, RTase, priming strategy and template concentration separately. Yield, as well as absolute copy numbers, were calculated as described in *Yield calculation* (see *Methods*).
